## Supplementary material for "On the Mathematics of RNA Velocity II: Algorithmic Aspects": Latex files: RNA.bib

%% This BibTeX bibliography file was created using BibDesk.
%% http://bibdesk.sourceforge.net/

%% Created for Tiejun Li at 2022-05-30 07:01:58 +0800 


%% Saved with string encoding Unicode (UTF-8) 


@article{mcinnes2018umap,
  title={{UMAP}: Uniform Manifold Approximation and Projection},
  author={McInnes, Leland and Healy, John and Saul, Nathaniel and Gro{\ss}berger, Lukas},
  journal={Journal of Open Source Software},
  volume={3},
  number={29},
  year={2018}
}

@article{Tang2009,
  title={m{RNA}-Seq whole-transcriptome analysis of a single cell},
  author={Tang, Fuchou and Barbacioru, Catalin and Wang, Yangzhou and Nordman, Ellen and Lee, Clarence and Xu, Nanlan and Wang, Xiaohui and Bodeau, John and Tuch, Brian B and Siddiqui, Asim and others},
  journal={Nature Methods},
  volume={6},
  number={5},
  pages={377--382},
  year={2009},
  publisher={Nature Publishing Group}
}

@article{MannoRNA2018,
  title={{RNA} velocity of single cells},
  author={La Manno, Gioele and Soldatov, Ruslan and Zeisel, Amit and Braun, Emelie and Hochgerner, Hannah and Petukhov, Viktor and Lidschreiber, Katja and Kastriti, Maria E and L{\"o}nnerberg, Peter and Furlan, Alessandro and others},
  journal={Nature},
  volume={560},
  number={7719},
  pages={494--498},
  year={2018},
  publisher={Nature Publishing Group}
}

@article{Weng2021,
  title={VeTra: a tool for trajectory inference based on RNA velocity},
  author={Weng, Guangzheng and Kim, Junil and Won, Kyoung Jae},
  journal={Bioinformatics},
  volume={37},
  number={20},
  pages={3509--3513},
  year={2021},
  publisher={Oxford University Press}
}

@article{Zhang2021,
  title={Inference of high-resolution trajectories in single-cell {RNA}-seq data by using {RNA} velocity},
  author={Zhang, Ziqi and Zhang, Xiuwei},
  journal={Cell Reports Methods},
  volume={1},
  number={6},
  pages={100095},
  year={2021},
  publisher={Elsevier}
}

@article{Atta2022,
  title={VeloViz: RNA velocity-informed embeddings for visualizing cellular trajectories},
  author={Atta, Lyla and Sahoo, Arpan and Fan, Jean},
  journal={Bioinformatics},
  volume={38},
  number={2},
  pages={391--396},
  year={2022},
  publisher={Oxford University Press}
}

@article{gorin2022rna,
  title={{RNA} velocity unraveled},
  author={Gorin, Gennady and Fang, Meichen and Chari, Tara and Pachter, Lior},
  journal={PLOS Computational Biology},
  volume={18},
  number={9},
  pages={e1010492},
  year={2022},
  publisher={Public Library of Science San Francisco, CA USA}
}

@article{Farrell2022,
  title={Inferring single-cell transcriptomic dynamics with structured dynamical representations of {RNA} velocity},
  author={Farrell, Spencer and Mani, Madhav and Goyal, Sidhartha},
  journal={Bulletin of the American Physical Society},
  year={2023},
  publisher={APS}
}

@article{Li1,
  title={On the mathematics of {RNA} Velocity {I}: theoretical analysis},
  author={Li, Tiejun and Shi, Jifan and Wu, Yichong and Zhou, Peijie},
  journal={CSIAM Transaction on Applied Mathematics},
  year = {2021},
  volume = {2},
  number = {1},
  pages = {1--55},
  publisher={Cold Spring Harbor Laboratory}
}

@article{spall2005monte,
  title={{Monte Carlo} computation of the {Fisher} information matrix in nonstandard settings},
  author={Spall, James C},
  journal={Journal of Computational and Graphical Statistics},
  volume={14},
  number={4},
  pages={889--909},
  year={2005},
  publisher={Taylor \& Francis}
}

@inproceedings{meng2017efficient,
  title={Efficient computation of the {Fisher} information matrix in the {EM} algorithm},
  author={Meng, Lingyao and Spall, James C},
  booktitle={2017 51st Annual Conference on Information Sciences and Systems (CISS)},
  pages={1--6},
  year={2017},
  organization={IEEE}
}

@article{MengUsing1991,
  title={Using {EM} to obtain asymptotic variance-covariance matrices: The {SEM} algorithm},
  author={Meng, Xiao-Li and Rubin, Donald B},
  journal={Journal of the American Statistical Association},
  volume={86},
  number={416},
  pages={899--909},
  year={1991},
  publisher={Taylor \& Francis}
}

@article{BergenGene2020,
  title={Generalizing {RNA} velocity to transient cell states through dynamical modeling},
  author={Bergen, Volker and Lange, Marius and Peidli, Stefan and Wolf, F Alexander and Theis, Fabian J},
  journal={Nature Biotechnology},
  volume={38},
  number={12},
  pages={1408--1414},
  year={2020},
  publisher={Nature Publishing Group}
}


@article{Oehlert1992,
  title={A note on the delta method},
  author={Oehlert, Gary W},
  journal={The American Statistician},
  volume={46},
  number={1},
  pages={27--29},
  year={1992},
  publisher={Taylor \& Francis}
}

@article{Singer,
  title={From graph to manifold Laplacian: The convergence rate},
  author={Singer, Amit},
  journal={Applied and Computational Harmonic Analysis},
  volume={21},
  number={1},
  pages={128--134},
  year={2006},
  publisher={Elsevier}
}


@article{Lange2022,
  title={CellRank for directed single-cell fate mapping},
  author={Lange, Marius and Bergen, Volker and Klein, Michal and Setty, Manu and Reuter, Bernhard and Bakhti, Mostafa and Lickert, Heiko and Ansari, Meshal and Schniering, Janine and Schiller, Herbert B and others},
  journal={Nature Methods},
  volume={19},
  number={2},
  pages={159--170},
  year={2022},
  publisher={Nature Publishing Group}
}


@article{Saelens2019,
  title={A comparison of single-cell trajectory inference methods},
  author={Saelens, Wouter and Cannoodt, Robrecht and Todorov, Helena and Saeys, Yvan},
  journal={Nature Biotechnology},
  volume={37},
  number={5},
  pages={547--554},
  year={2019},
  publisher={Nature Publishing Group}
}

@article{Trapnell2014,
  title={The dynamics and regulators of cell fate decisions are revealed by pseudotemporal ordering of single cells},
  author={Trapnell, Cole and Cacchiarelli, Davide and Grimsby, Jonna and Pokharel, Prapti and Li, Shuqiang and Morse, Michael and Lennon, Niall J and Livak, Kenneth J and Mikkelsen, Tarjei S and Rinn, John L},
  journal={Nature Biotechnology},
  volume={32},
  number={4},
  pages={381--386},
  year={2014},
  publisher={Nature Publishing Group}
}


@article{gao2022unitvelo,
  title={{UniTVelo}: temporally unified {RNA} velocity reinforces single-cell trajectory inference},
  author={Gao, Mingze and Qiao, Chen and Huang, Yuanhua},
  journal={Nature Communications},
   volume={13},
  number={1},
  pages={6586},
  year={2022},
  publisher={Cold Spring Harbor Laboratory}
}

@article{marot2022towards,
  title={Towards reliable quantification of cell state velocities},
  author={Marot-Lassauzaie, Val{\'e}rie and Bouman, Brigitte Joanne and Donaghy, Fearghal Declan and Demerdash, Yasmin and Essers, Marieke Alida Gertruda and Haghverdi, Laleh},
  journal={PLoS Computational Biology},
  volume={18},
  number={9},
  pages={e1010031},
  year={2022},
  publisher={Public Library of Science San Francisco, CA USA}
}

@article{gayoso2022deep,
  title={Deep generative modeling of transcriptional dynamics for {RNA} velocity analysis in single cells},
  author={Gayoso, Adam and Weiler, Philipp and Lotfollahi, Mohammad and Klein, Dominik and Hong, Justin and Streets, Aaron M and Theis, Fabian J and Yosef, Nir},
  journal={bioRxiv},
  year={2022},
  publisher={Cold Spring Harbor Laboratory}
}

@article{qin2022pyro,
  title={{Pyro-Velocity}: Probabilistic {RNA} Velocity inference from single-cell data},
  author={Qin, Qian and Bingham, Eli and La Manno, Gioele and Langenau, David M and Pinello, Luca},
  journal={bioRxiv},
  year={2022},
  publisher={Cold Spring Harbor Laboratory}
}

@article{weinreb2018fundamental,
  title={Fundamental limits on dynamic inference from single-cell snapshots},
  author={Weinreb, Caleb and Wolock, Samuel and Tusi, Betsabeh K and Socolovsky, Merav and Klein, Allon M},
  journal={Proceedings of the National Academy of Sciences},
  volume={115},
  number={10},
  pages={E2467--E2476},
  year={2018},
  publisher={National Acad Sciences}
}

@article{zhou2021dissecting,
  title={Dissecting transition cells from single-cell transcriptome data through multiscale stochastic dynamics},
  author={Zhou, Peijie and Wang, Shuxiong and Li, Tiejun and Nie, Qing},
  journal={Nature Communications},
  volume={12},
  number={1},
  pages={5609},
  year={2021},
  publisher={Nature Publishing Group UK London}
}

@article{setty2019characterization,
  title={Characterization of cell fate probabilities in single-cell data with Palantir},
  author={Setty, Manu and Kiseliovas, Vaidotas and Levine, Jacob and Gayoso, Adam and Mazutis, Linas and Pe’Er, Dana},
  journal={Nature Biotechnology},
  volume={37},
  number={4},
  pages={451--460},
  year={2019},
  publisher={Nature Publishing Group US New York}
}

@article{townes2019feature,
  title={Feature selection and dimension reduction for single-cell RNA-Seq based on a multinomial model},
  author={Townes, F William and Hicks, Stephanie C and Aryee, Martin J and Irizarry, Rafael A},
  journal={Genome biology},
  volume={20},
  pages={1--16},
  year={2019},
  publisher={Springer}
}

@article{luecken2019current,
  title={Current best practices in single-cell RNA-seq analysis: a tutorial},
  author={Luecken, Malte D and Theis, Fabian J},
  journal={Molecular systems biology},
  volume={15},
  number={6},
  pages={e8746},
  year={2019}
}

@article{qiao2021representation,
  title={Representation learning of {RNA} velocity reveals robust cell transitions},
  author={Qiao, Chen and Huang, Yuanhua},
  journal={Proceedings of the National Academy of Sciences},
  volume={118},
  number={49},
  pages={e2105859118},
  year={2021},
  publisher={National Acad Sciences}
}

@article{atta2022veloviz,
  title={VeloViz: {RNA} velocity-informed embeddings for visualizing cellular trajectories},
  author={Atta, Lyla and Sahoo, Arpan and Fan, Jean},
  journal={Bioinformatics},
  volume={38},
  number={2},
  pages={391--396},
  year={2022},
  publisher={Oxford Academic}
}


@article{liu2022dynamical,
  title={Dynamical systems model of {RNA} velocity improves inference of single-cell trajectory, pseudo-time and gene regulation},
  author={Liu, Ruishan and Pisco, Angela Oliveira and Braun, Emelie and Linnarsson, Sten and Zou, James},
  journal={Journal of Molecular Biology},
  volume={434},
  number={15},
  pages={167606},
  year={2022},
  publisher={Elsevier}
}

@book{norris1998markov,
  title={Markov Chains},
  author={Norris, James R},
  year={1998},
  publisher={Cambridge University Press},
  address={Cambridge}
}


@article{eling2019challenges,
  title={Challenges in measuring and understanding biological noise},
  author={Eling, Nils and Morgan, Michael D and Marioni, John C},
  journal={Nature Reviews Genetics},
  volume={20},
  number={9},
  pages={536--548},
  year={2019},
  publisher={Nature Publishing Group UK London}
}

@article{hilfinger2011separating,
  title={Separating intrinsic from extrinsic fluctuations in dynamic biological systems},
  author={Hilfinger, Andreas and Paulsson, Johan},
  journal={Proceedings of the National Academy of Sciences},
  volume={108},
  number={29},
  pages={12167--12172},
  year={2011},
  publisher={National Acad Sciences}
}

@article{zhou2021stochasticity,
  title={Stochasticity triggers activation of the {S-phase} checkpoint pathway in budding yeast},
  author={Zhou, Peijie and Gao, Xin and Li, Xiaoli and Li, Linxi and Niu, Caoyuan and Ouyang, Qi and Lou, Huiqiang and Li, Tiejun and Li, Fangting},
  journal={Physical Review X},
  volume={11},
  number={1},
  pages={011004},
  year={2021},
  publisher={APS}
}

@article{qiu2022mapping,
  title={Mapping transcriptomic vector fields of single cells},
  author={Qiu, Xiaojie and Zhang, Yan and Martin-Rufino, Jorge D and Weng, Chen and Hosseinzadeh, Shayan and Yang, Dian and Pogson, Angela N and Hein, Marco Y and Min, Kyung Hoi Joseph and Wang, Li and others},
  journal={Cell},
  volume={185},
  number={4},
  pages={690--711},
  year={2022},
  publisher={Elsevier}
}


@article{Schur1911lemma1,
url = {https://doi.org/10.1515/crll.1911.140.1},
title = {Bemerkungen zur Theorie der beschränkten Bilinearformen mit unendlich vielen Veränderlichen.},
title = {},
author = {J. Schur},
pages = {1--28},
volume = {1911},
number = {140},
journal = {},
doi = {doi:10.1515/crll.1911.140.1},
year = {1911},
lastchecked = {2023-04-14}
}

@book{Horn85,
title={Matrix Analysis},
author={Horn, R.A. and Johnson, C.R.},
year={1985},
publisher={Cambridge University Press},
address={Cambridge}
}

@unpublished{Wang23Preprint,
title = {Inferring the {RNA} velocity with stochastic models},
author = {Wang, Yizhuo and Li, Tiejun},
note={preprint},
year = {2023}
}

@article{trapnell2014pseudo,
  title={Pseudo-temporal ordering of individual cells reveals dynamics and regulators of cell fate decisions},
  author={Trapnell, Cole and Cacchiarelli, Davide and Grimsby, Jonna and Pokharel, Prapti and Li, Shuqiang and Morse, Michael and Lennon, Niall J and Livak, Kenneth J and Mikkelsen, Tarjei S and Rinn, John L},
  journal={Nature Biotechnology},
  volume={32},
  number={4},
  pages={381},
  year={2014},
  publisher={NIH Public Access}
}

@article{haghverdi2016diffusion,
  title={Diffusion pseudotime robustly reconstructs lineage branching},
  author={Haghverdi, Laleh and B{\"u}ttner, Maren and Wolf, F Alexander and Buettner, Florian and Theis, Fabian J},
  journal={Nature Methods},
  volume={13},
  number={10},
  pages={845--848},
  year={2016},
  publisher={Nature Publishing Group US New York}
}

@article{wolf2019paga,
  title={{PAGA}: graph abstraction reconciles clustering with trajectory inference through a topology preserving map of single cells},
  author={Wolf, F Alexander and Hamey, Fiona K and Plass, Mireya and Solana, Jordi and Dahlin, Joakim S and G{\"o}ttgens, Berthold and Rajewsky, Nikolaus and Simon, Lukas and Theis, Fabian J},
  journal={Genome Biology},
  volume={20},
  pages={1--9},
  year={2019},
  publisher={Springer}
}

@article{von2014hitting,
  title={Hitting and commute times in large random neighborhood graphs},
  author={Von Luxburg, Ulrike and Radl, Agnes and Hein, Matthias},
  journal={The Journal of Machine Learning Research},
  volume={15},
  number={1},
  pages={1751--1798},
  year={2014},
  publisher={JMLR. org}
}

@article{stassen2021generalized,
  title={Generalized and scalable trajectory inference in single-cell omics data with {VIA}},
  author={Stassen, Shobana V and Yip, Gwinky GK and Wong, Kenneth KY and Ho, Joshua WK and Tsia, Kevin K},
  journal={Nature Communications},
  volume={12},
  number={1},
  pages={5528},
  year={2021},
  publisher={Nature Publishing Group UK London}
}

@article{bocci2022splicejac,
  title={SpliceJAC: transition genes and state-specific gene regulation from single-cell transcriptome data},
  author={Bocci, Federico and Zhou, Peijie and Nie, Qing},
  journal={Molecular Systems Biology},
  volume={18},
  number={11},
  pages={e11176},
  year={2022}
}

@article{wang2022inference,
  title={Inference on the structure of gene regulatory networks},
  author={Wang, Yue and Wang, Zikun},
  journal={Journal of Theoretical Biology},
  volume={539},
  pages={111055},
  year={2022},
  publisher={Elsevier}
}

@article{li2023multi,
  title={Multi-omic single-cell velocity models epigenome--transcriptome interactions and improves cell fate prediction},
  author={Li, Chen and Virgilio, Maria C and Collins, Kathleen L and Welch, Joshua D},
  journal={Nature Biotechnology},
  volume={41},
  number={3},
  pages={387--398},
  year={2023},
  publisher={Nature Publishing Group US New York}
}

@article{gu2022bayesian,
  title={Bayesian inference of {RNA} velocity from multi-lineage single-cell data},
  author={Gu, Yichen and Blaauw, David and Welch, Joshua D},
  journal={bioRxiv},
  pages={2022--07},
  year={2022},
  publisher={Cold Spring Harbor Laboratory}
}

@book{Lehmann1998,
  title={Theory of Point Estimation},
  author={Lehmann, E.L. and Casella, G.},
  year={1998},
  publisher={Springer},
  address={New York},
  edition={2nd}
}

@article{hashimoto2015random,
  title={From random walks to distances on unweighted graphs},
  author={Hashimoto, Tatsunori and Sun, Yi and Jaakkola, Tommi},
  journal={Advances in Neural Information Processing Systems},
  volume={28},
  year={2015}
}

@article{gorin2022interpretable,
  title={Interpretable and tractable models of transcriptional noise for the rational design of single-molecule quantification experiments},
  author={Gorin, Gennady and Vastola, John J and Fang, Meichen and Pachter, Lior},
  journal={Nature Communications},
  volume={13},
  number={1},
  pages={7620},
  year={2022},
  publisher={Nature Publishing Group UK London}
}

@book{Kemeny76,
	address = {New York},
	author = {Kemeny, J.G. and Snell, J.L.},
	date-added = {2023-05-15 11:00:40 +0800},
	date-modified = {2023-05-15 11:01:50 +0800},
	edition = {2nd},
	publisher = {Springer},
	title = {Finite Markov Chains},
	year = {1976}}

@article{ma2013regularized,
  title={Regularized vector field learning with sparse approximation for mismatch removal},
  author={Ma, Jiayi and Zhao, Ji and Tian, Jinwen and Bai, Xiang and Tu, Zhuowen},
  journal={Pattern Recognition},
  volume={46},
  number={12},
  pages={3519--3532},
  year={2013},
  publisher={Elsevier}
}

@article{chen2022deepvelo,
  title={{DeepVelo}: Single-cell transcriptomic deep velocity field learning with neural ordinary differential equations},
  author={Chen, Zhanlin and King, William C and Hwang, Aheyon and Gerstein, Mark and Zhang, Jing},
  journal={Science Advances},
  volume={8},
  number={48},
  pages={eabq3745},
  year={2022},
  publisher={American Association for the Advancement of Science}
}

@article{chen2018neural,
  title={Neural ordinary differential equations},
  author={Chen, Ricky TQ and Rubanova, Yulia and Bettencourt, Jesse and Duvenaud, David K},
  journal={Advances in Neural Information Processing Systems},
  volume={31},
  year={2018}
}

@article{gorin2022modeling,
  title={Modeling bursty transcription and splicing with the chemical master equation},
  author={Gorin, Gennady and Pachter, Lior},
  journal={Biophysical Journal},
  volume={121},
  number={6},
  pages={1056--1069},
  year={2022},
  publisher={Elsevier}
}

@article {Zhang2023learning,
	author = {Stephen Y Zhang and Michael P H Stumpf},
	title = {Learning cell-specific networks from dynamical single cell data},
	year = {2023},
	doi = {10.1101/2023.01.08.523176},
	publisher = {Cold Spring Harbor Laboratory},
	journal = {bioRxiv}
}

@article{dong2022integrating,
  title={Integrating single-cell datasets with ambiguous batch information by incorporating molecular network features},
  author={Dong, Ji and Zhou, Peijie and Wu, Yichong and Chen, Yidong and Xie, Haoling and Gao, Yuan and Lu, Jiansen and Yang, Jingwei and Zhang, Xiannian and Wen, Lu and Li, Tiejun and Tang, Fuchou},
  journal={Briefings in Bioinformatics},
  volume={23},
  number={1},
  pages={bbab366},
  year={2022},
  month = {09},
  publisher={Oxford University Press}
}

@article{spigler2020asymptotic,
  title={Asymptotic learning curves of kernel methods: empirical data versus teacher--student paradigm},
  author={Spigler, Stefano and Geiger, Mario and Wyart, Matthieu},
  journal={Journal of Statistical Mechanics: Theory and Experiment},
  volume={2020},
  number={12},
  pages={124001},
  year={2020},
  publisher={IOP Publishing}
}
