## Supplementary figures and images for "On the Mathematics of RNA Velocity II: Algorithmic Aspects"

### CI-eps-converted-to.pdf

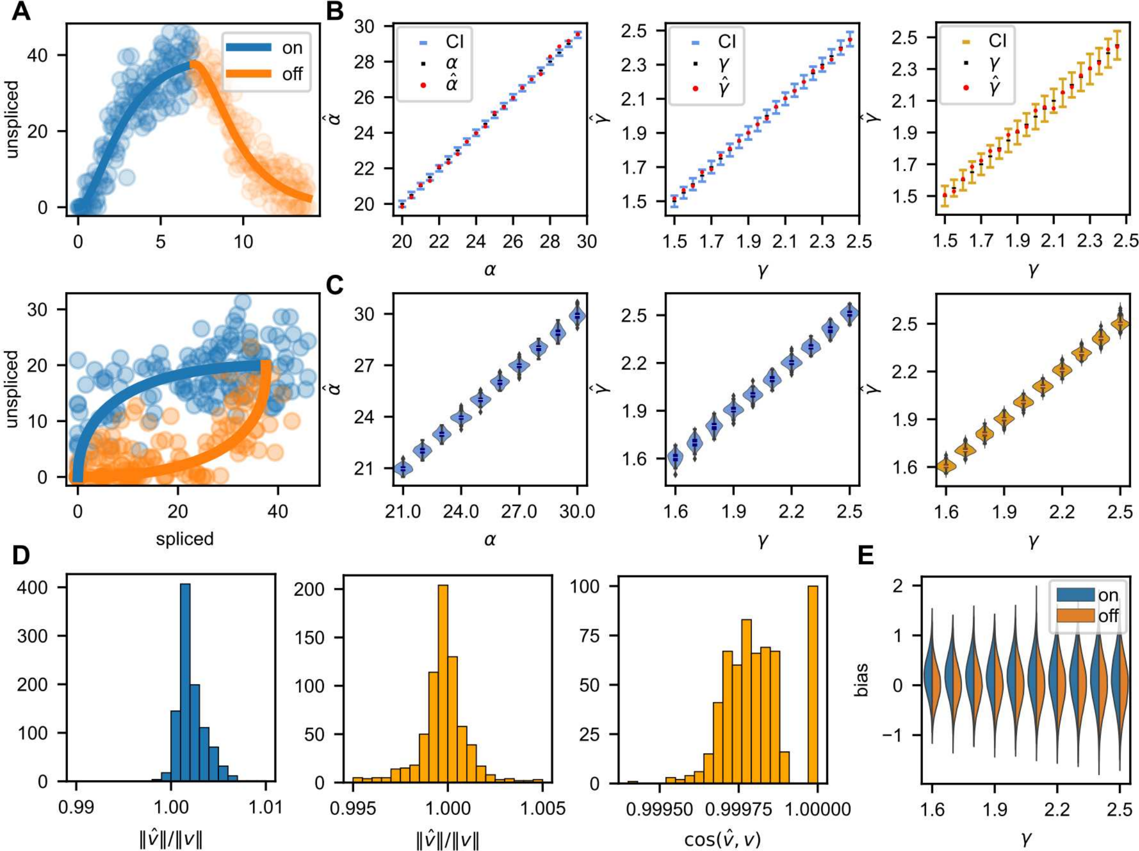

### fig1-eps-converted-to.pdf

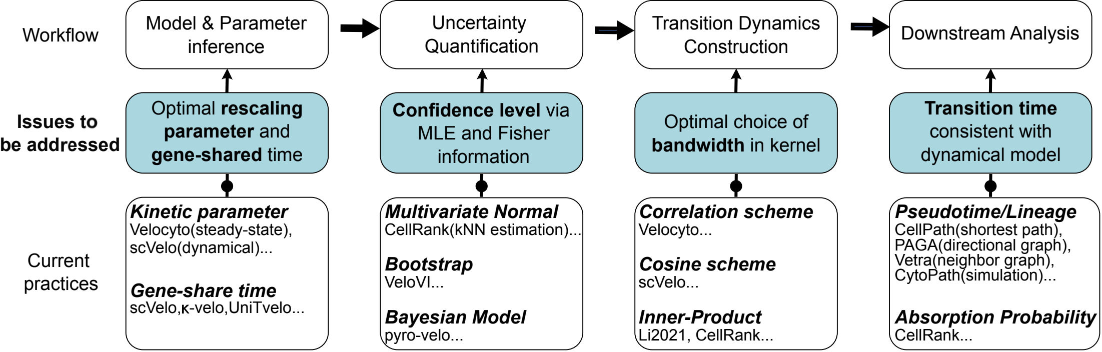

### fig4-eps-converted-to.pdf

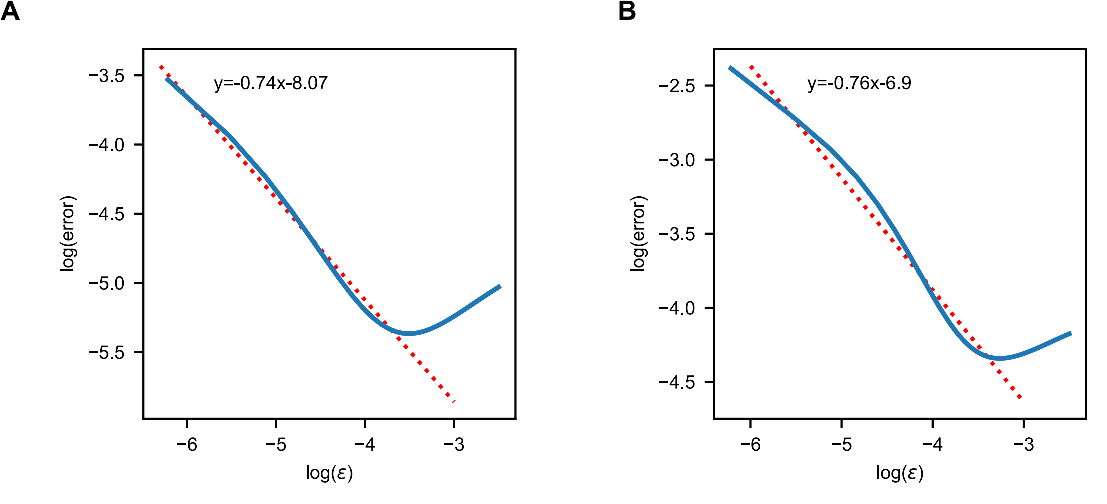

### fig5_v4-eps-converted-to.pdf

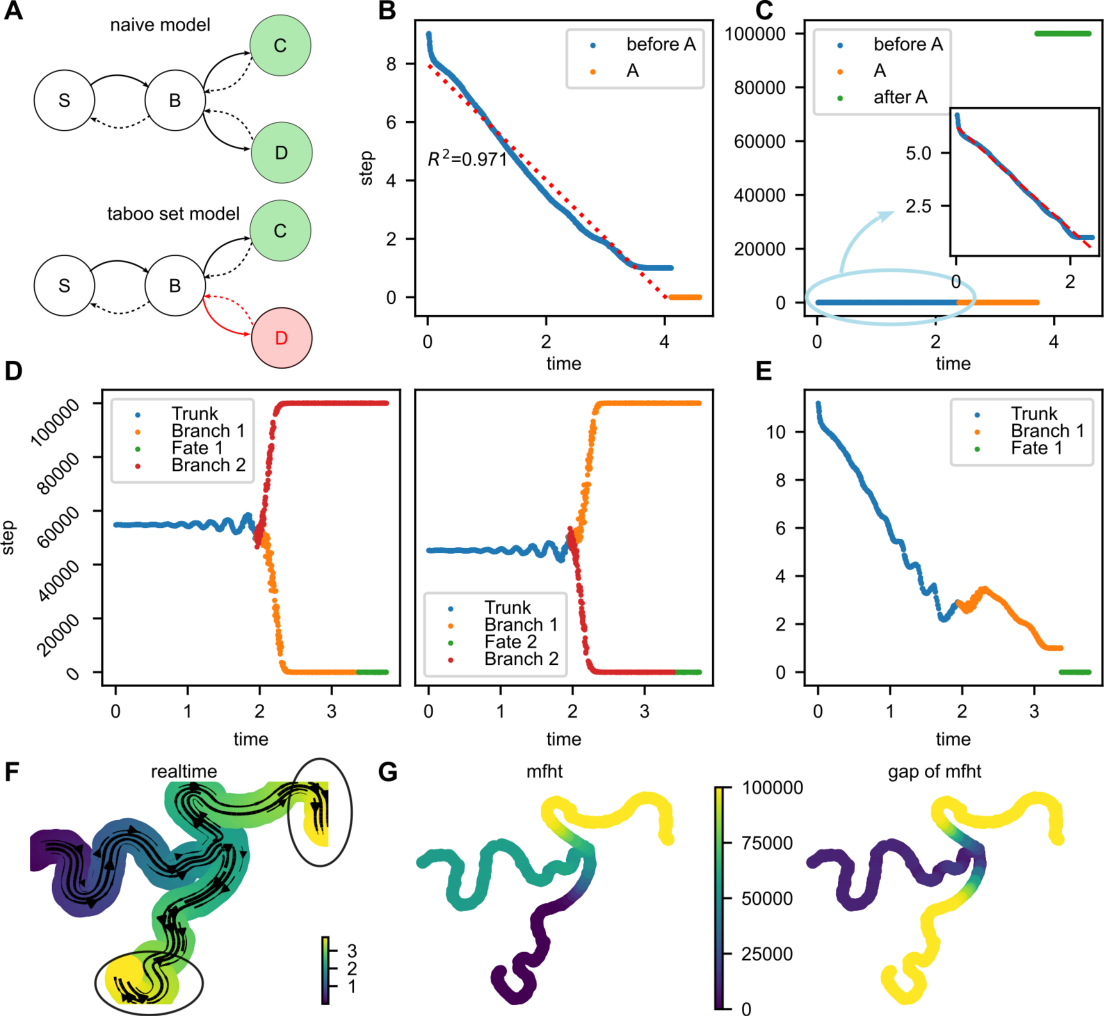

### rescale.pdf

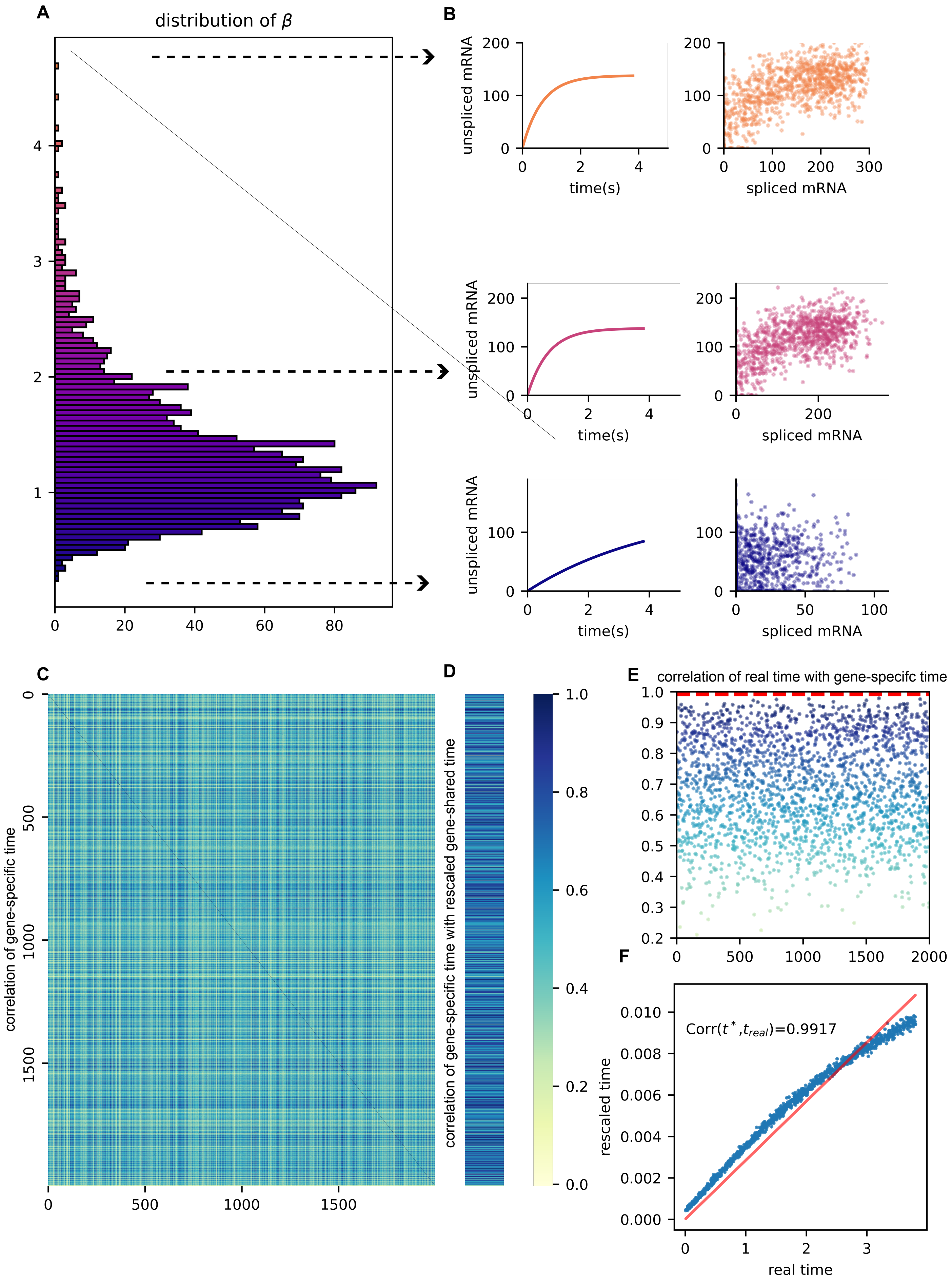

### rescale_v2.pdf

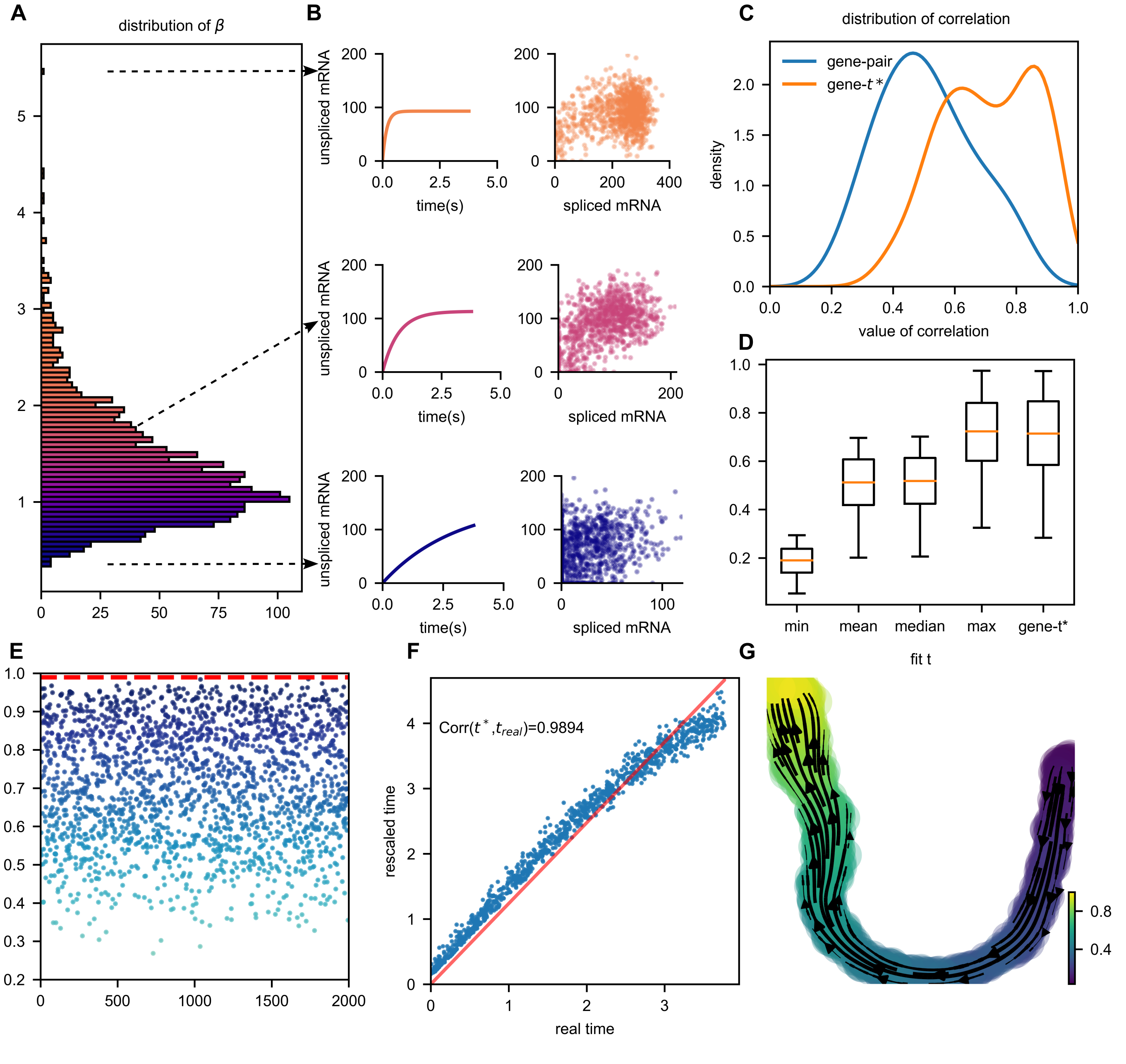

### rescale_v4-eps-converted-to.pdf

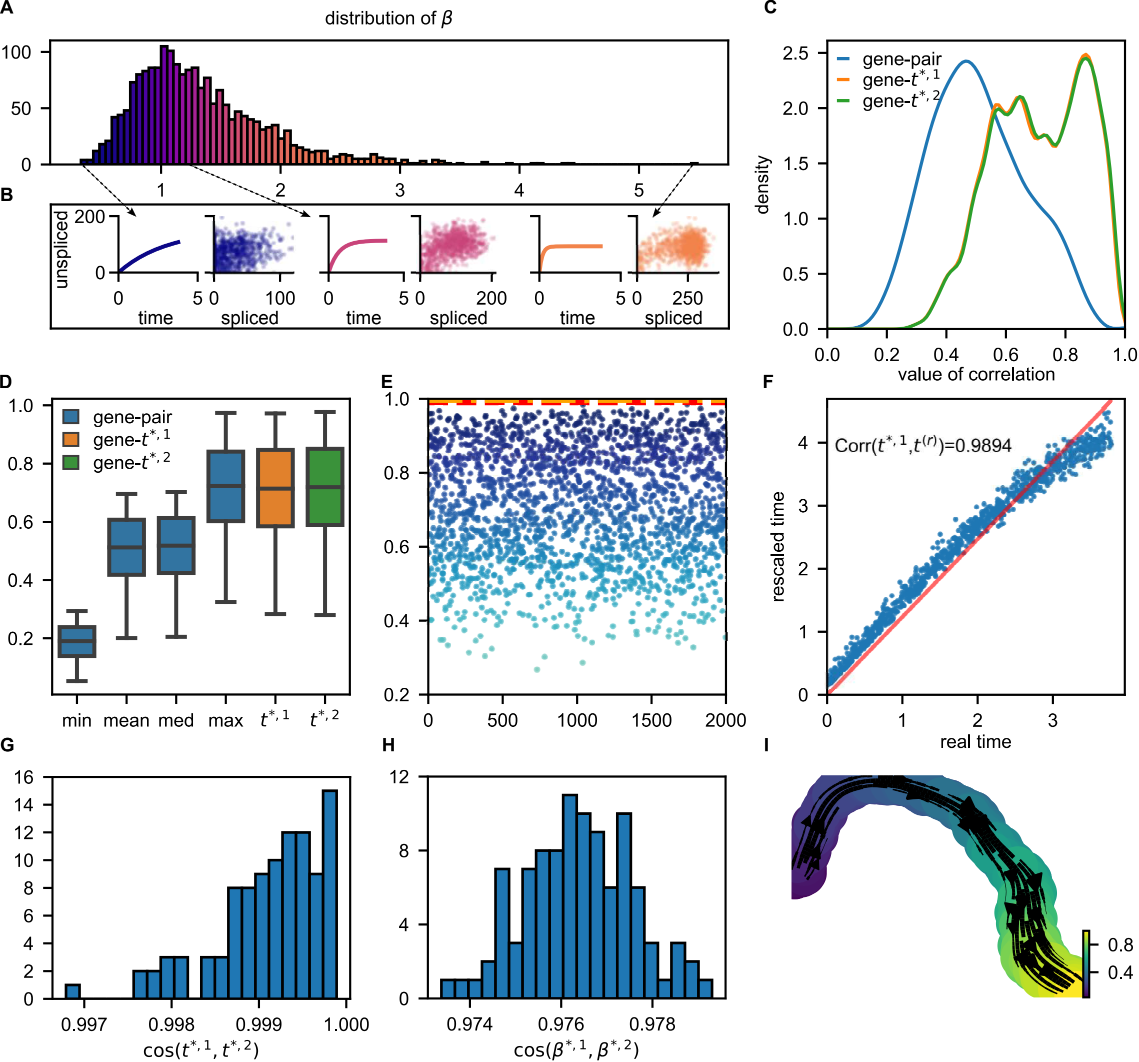
